# Optimisation strategies for directed evolution without sequencing

**DOI:** 10.1101/2024.03.18.585521

**Authors:** Jess James, Sebastian Towers, Jakob Foerster, Harrison Steel

## Abstract

Directed evolution can enable engineering of biological systems with minimal knowledge of their underlying sequence-to-function relationships. A typical directed evolution process consists of iterative rounds of mutagenesis and selection that are designed to steer changes in a biological system (e.g. a protein) towards some functional goal. Much work has been done, particularly leveraging advancements in machine learning, to optimise the process of directed evolution. Many of these methods, however, require DNA sequencing and synthesis, making them resource-intensive and incompatible with developments in targeted *in vivo* mutagenesis. Operating within the experimental constraints of established sorting-based directed evolution techniques (e.g. Fluorescence-Activated Cell Sorting, FACS), we explore approaches for optimisation of directed evolution that do not require sequencing information. We then expand our methods to the context of emerging experimental techniques in directed evolution, which allow for single-cell selection based on fitness objectives defined from any combination measurable traits. Finally, we validate the developed selection strategies on the GB1 empirical landscape, demonstrating that they can lead to up to a 7.5 fold increase in the probability of attaining the global fitness peak.

**Author summary:** The standard approach to sorting-based selection in directed evolution is to take forward only the top-performing variants from each generation of a single population. In this work, we begin to explore alternative selection strategies within a simulated directed evolution framework. We propose “selection functions”, which allow us to tune the balance of exploration and exploitation of a fitness landscape, and we demonstrate that splitting a population into sub-populations can improve both the likelihood and magnitude of a successful outcome. We also propose strategies to leverage emerging selection methods that can implement single-cell selection based on any combination of measurable traits. We validate our optimised directed evolution approaches on the empirical fitness landscape of the GB1 immunoglobulin protein.

## Introduction

Engineered biological systems hold immense potential for application across industries including medicine, manufacturing, and agriculture [1–3]. In recent decades, protein engineering in particular has demonstrated the potential of natural biological elements to be adapted for new functionalities. Advancements in computational methods are bringing *de novo* protein engineering closer to reality [4–8]. However, such approaches remain limited by our developing understanding of protein sequence-to-function relationships. As one means to circumvent the need for such detailed prior knowledge, a range of techniques termed “directed evolution” have been developed [9]. Directed evolution techniques have delivered products across a range of applications, from cancer and autoimmune disorder drugs [10] to enzymes for converting cooking oil into bio diesel [11].

In a process that mimics nature, directed evolution consists of iteratively introducing random variation by mutagenesis, followed by selection biased toward user-defined desirable variants. Selection can be achieved broadly via two families of approaches; first, those that couple a trait-of-interest to growth of a host organism [12, 13], or second, those that couple a trait to a measurable output (e.g. expression of fluorescent proteins) and then actively sort cells with desirable output values for future mutagenesis and propagation [14]. Each approach has different strengths and weaknesses. Growth-coupled selection utilises (comparatively) straightforward growth-based assays but requires engineering of a trait-to-growth coupling, which is both challenging and can lead to “cheating” behaviour [15]. Meanwhile, sorting-based methods only require a trait be *measurable* (i.e. they do not require coupling to growth), but in turn necessitate more complex experimental approaches to implement the selection for the measured winners (e.g. FACS [14]). In contrast to FACS, which takes only a single time-point measurement from each cell, emerging selection techniques leverage microfluidics to observe cells over long time periods prior to sorting [16, 17]. This produces increasingly high-dimensional information on which to sort cells. Our work focuses on these emerging sorting-based methods due to their increased level of control, and consequently not all strategies we propose are suitable for growth-coupled directed evolution. This is a timely challenge, as new methods for single-cell selection pose novel theoretical questions - which our work aims to answer - regarding how their capabilities can be optimally implemented and exploited.

As a trait-of-interest undergoes directed evolution, the process can be imagined as navigation across high-dimensional “fitness landscapes” [18]. Fitness landscapes map each genetic sequence to a measure of fitness, with “fitness” being performance for a desired function - the goal of directed evolution is to find the highest peaks on that landscape. Fitness landscapes are known to exhibit variable degrees of ruggedness, which can create local optima that constrain paths of evolution [19, 20]. Standard practice in directed evolution is to take forward and mutate only the top fraction of variants with each iteration [21, 22]. This “greedy” approach is prone to getting trapped in local optima, particularly in rugged landscapes [23]. With the aid of computational methods, however, it is possible to navigate protein fitness landscapes in a more active way. One of the earliest examples of such a method is ProSAR, which uses a statistical algorithm to identify specific residues that are correlated with high fitness. Each new generation of variants is designed to combine residues that were predicted to contribute most to fitness [24]. Methods that predict the fitness effects of mutations in this way are now able to accommodate machine learning [25–27] and Bayesian optimisation approaches [28, 29]. Such methods have been built upon by not only utilising the fitnesses of the sequences in isolation, but also time-series mutation data acquired *during* a directed evolution experiment [30]. The vast majority of these methods, however, are based upon the requirement for sequencing information from each generation. This means they are somewhat resource- and labour-intensive, and are not suited to maximising the benefits from *in vivo* mutagenesis methods for directed evolution [31].

Here we explore how, in the absence of sequencing data, one can maximise the likelihood of a given directed evolution process finding global (or at least very strong local) optima on a fitness landscape. Examples of previous work to optimise directed evolution without sequencing information include strategies such as alternating between “on” and “off” states of selection [32]. This approach offers an opportunity for populations to traverse fitness valleys and avoid getting trapped at local optima. Here, we approach the same challenge from several new angles. First, probability of selection is applied as a parameterised function of fitness that can be used to tune the balance exploration and exploitation on a fitness landscape. Second, we investigate the benefits that can be gained by splitting a population into sub-populations and allowing their trajectories to diverge. Finally, we explore the novel capabilities of the aforementioned emerging selection methods [16, 17], which in particular enable effective optimisation of multiple properties in parallel. We demonstrate the performance of our optimisation approaches by simulating directed evolution on the GB1 empirical landscape [19].

## Results

In order to test selection strategies, a computational model was implemented to mimic the process of directed evolution. Genes in the model are represented by one-dimensional arrays, which iterate through rounds of mutation and selection (Fig. 1, Methods: Model). In the selection process, the fitness of each gene is calculated using an empirical landscape [19] or an NK model [33, 34]; see *Methods*. Empirical landscapes are combinatorially complete fitness measurements for all variants of a protein (or protein region). NK models are computationally generated fitness landscapes that are made necessary by the limited availability of empirical landscapes. In both NK and empirical landscape implementations, the gene sequence can be taken as input and the corresponding fitness value is given as output. As outlined above, the approaches explored with the model are possible in contexts where one actively selects “winning” variants to enrich (e.g. FACS), as opposed to growth-coupled directed evolution, which does not offer this type of control.

**Fig 1.**
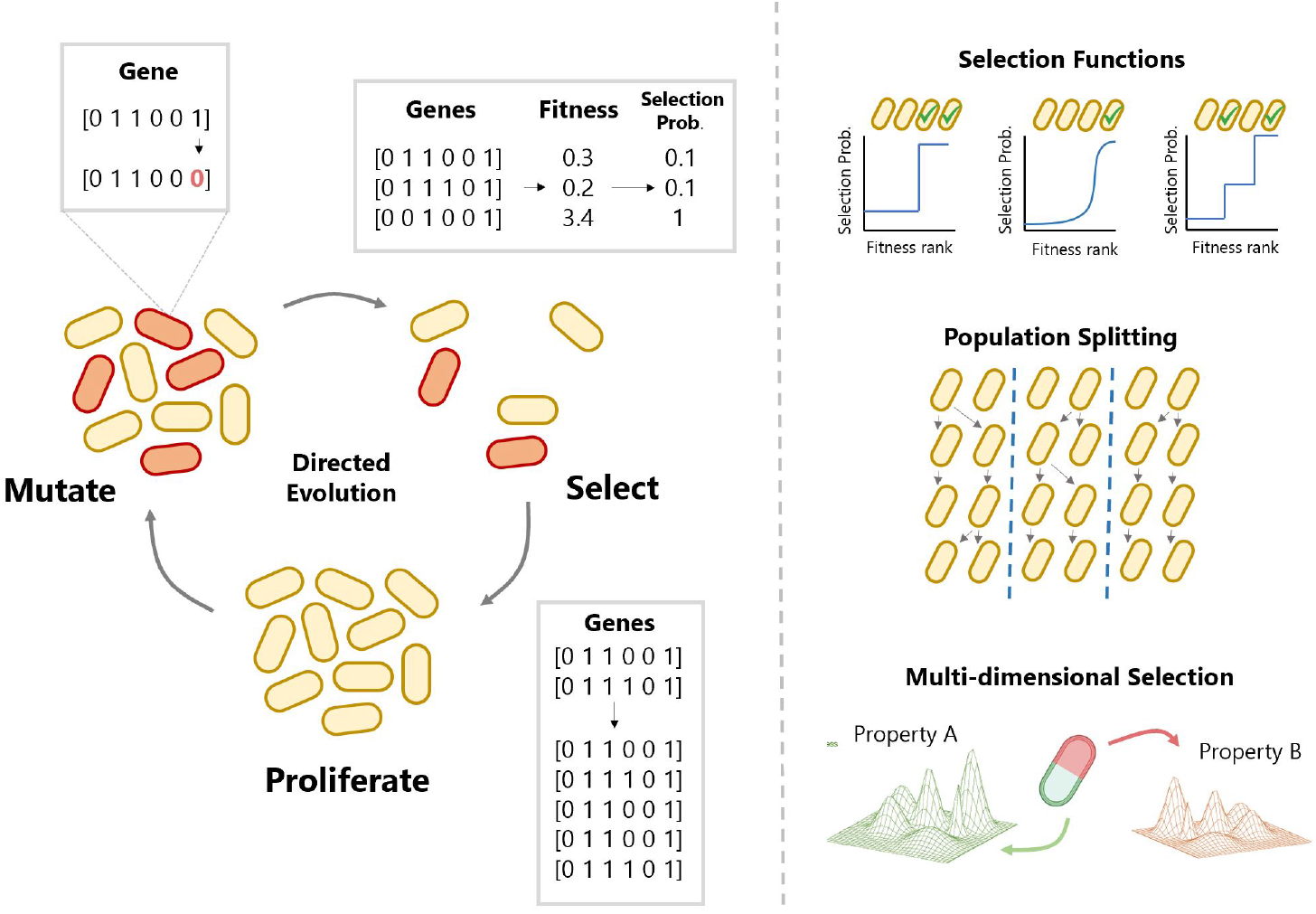
Schematic of the directed evolution simulation cycle. The model of directed evolution performs iterative round of mutation, selection and proliferation. Genes are represented by one-dimensional arrays. The fitness of each gene can be generated by feeding the array into a fitness landscape model. The probability of selection is determined by feeding the resulting fitness into a selection function. Proliferation is carried out by sampling with even probability up to a fixed population size. Mutation is carried out by introducing random changes to the arrays. For a more detailed description of the computational pipeline, see Methods: Model. Strategies explored using the model include selection functions, population splitting and selection across multiple properties

### Selection functions for tuneable exploration vs exploitation

Selection functions are introduced as a means to tune the balance of exploration and exploitation on a fitness landscape. The selection functions proposed here are defined by two parameters: “threshold” and “base chance” (Fig. 2A). The threshold is the fitness percentile above which variants have a 100% chance of selection, otherwise the chance of selection is equal to the base chance (Methods: Selection Functions). In this work, selection functions are normalised to select a constant fraction of variants. In a continuous directed evolution experiment, this ensures that proliferation time remains approximately constant between generations (hence, performance metrics can be considered as improvement in a trait *per unit time*). Fixing the selected proportion of cells also reduces the parameter space of selection functions to one dimension, as every base chance has only one threshold value corresponding to a fixed proportion of the population being selected. We hypothesise that the base chance parameter will improve directed evolution by allowing a population to escape local optima on the fitness landscape. By selecting some cells unconditionally, they are allowed to accumulate more mutations, potentially allowing them reach higher performing variants via deleterious phenotypes.

**Fig 2.**
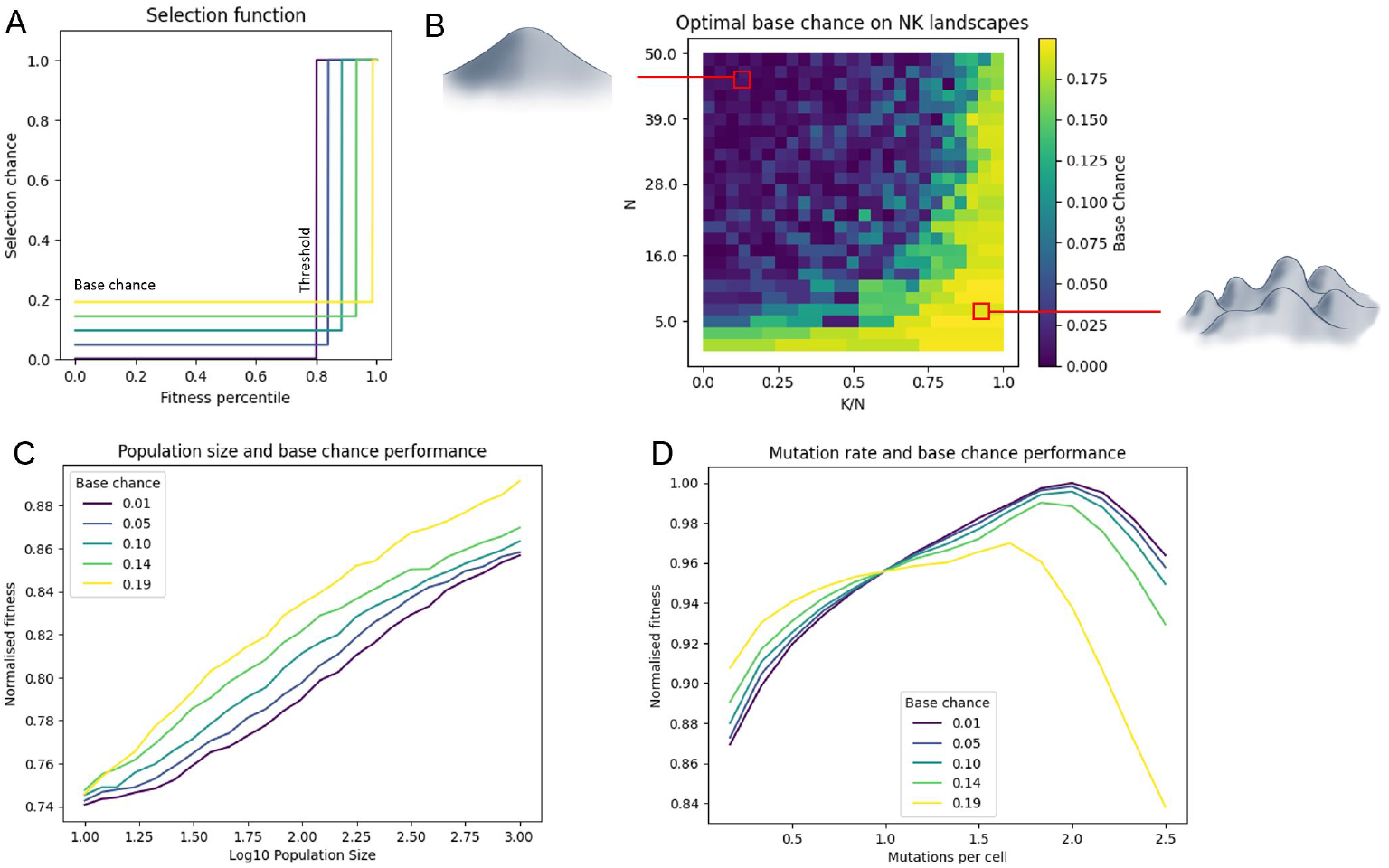
Investigating the performance of selection functions in directed evolution. A: Selection functions define probability of selection as a function of fitness (Methods : Selection Functions). The selection function used here is determined by two parameters: threshold and base chance. Selection functions were normalised to select 20% of the population. B: Optimal base chance values on varying NK landscapes. Dependence of trajectory end-point fitness on C: mutation rate, and D: population size. “Normalised fitness” is maximum fitness across the population at the final time point (averaged over 100 runs) as a fraction of the global maximum on the landscape. Experiments ran for 300 generations. 0.01 *≤* base chance *≤* 0.19, N = 25, K = 5, mutations per cell = 0.1, population size = 1000.

The NK model describes a class of fitness landscape with tuneable ruggedness [33, 34] (Methods: NK Landscapes). N describes the number of variable sites, and K can be thought of as a metric of ruggedness ranging from 0 to *N* − 1, with high K meaning high ruggedness (i.e. more local optima). Fig. 2B shows the optimal base chance over varying NK landscapes, as estimated via simulation. In particular, Fig. 2B shows the optimal base chance increasing with respect to *K* (ruggedness), and decreasing with respect to *N* (dimensionality). Given that base chance is hypothesised to help escape local optima, one explanation for this result is that more local optima are found in rugged landscapes, and they are more difficult to escape in low-dimensional landscapes, which offer fewer paths between any two points. For smooth, high-dimensional landscapes the opposite is true, therefore the function that most favours exploitation over exploration (i.e. base chance = 0) is found to be optimal.

Next, the interaction between base chance and population size was explored. Note that the NK landscape is non-linear, therefore a 1% increase in raw fitness may truly represent a larger underlying improvement, particularly at the high-fitness end of the distribution Fig. 2C shows that base chance can improve performance across a range of population sizes, particularly in large populations. Given that large populations can explore more of the landscape, they are likely to encounter more local optima. The ability for a population to escape a local optimum is dependent on its ability to reach a fitter state via at least one deleterious mutation. Although evolution via deleterious mutations is shown to occur more readily in large populations [35, 36], the dynamic is also promoted by base chance, and hence base chance can improve performance. In small populations, the cost of including detrimental variants is greater relative to the potential gain, therefore base chance is less beneficial.

The performance of the selection functions with varying mutation rates was also investigated (Fig. 2D). When mutation rate is low, higher base chance values perform best and vice versa for high mutation rate. One explanation for this is in the balance of exploration and exploitation. Both base chance and mutation rate aid in escaping local optima by increasing the likelihood of a cell undergoing multiple mutations. Although this benefits exploration, *further* increases in mutation rate can come at the cost of not effectively exploiting a position on the landscape. For this reason, base chance performance drops off more quickly in a high mutation rate regime. Given that most directed evolution experiments operate in a low mutation rate regime to avoid detrimental side effects [37], base chance could act as a useful tool for promoting landscape exploration. The benefits of such an approach are that implementing a base chance has no direct impact on top-performing variants, whereas increasing mutation rate impacts all cells.

### Population splitting for improved exploration

Until now, this work has assumed a single population undergoing directed evolution. However, in practice, one could run multiple, smaller, copies of the same directed evolution experiment by subdividing a population, and take the best outcome across all of them as the final result. Here, this method is referred to as “population splitting”. An example of such a situation is displayed in Fig.3A, where a population of size 500 is split into five equal sub-populations of size 100. In this example experiment population splitting performs better, and Fig. 3D and E demonstrate the consistency of this result across parameter regimes. This may be because in a single, mixed population, mutations will eventually drift to fixation or extinction, therefore the population as a whole remains largely on the same trajectory. If one splits the population, sub-populations are able to drift on separate trajectories without cross-competition, effectively mimicking the process of speciation and increasing landscape exploration [38, 39].

**Fig 3.**
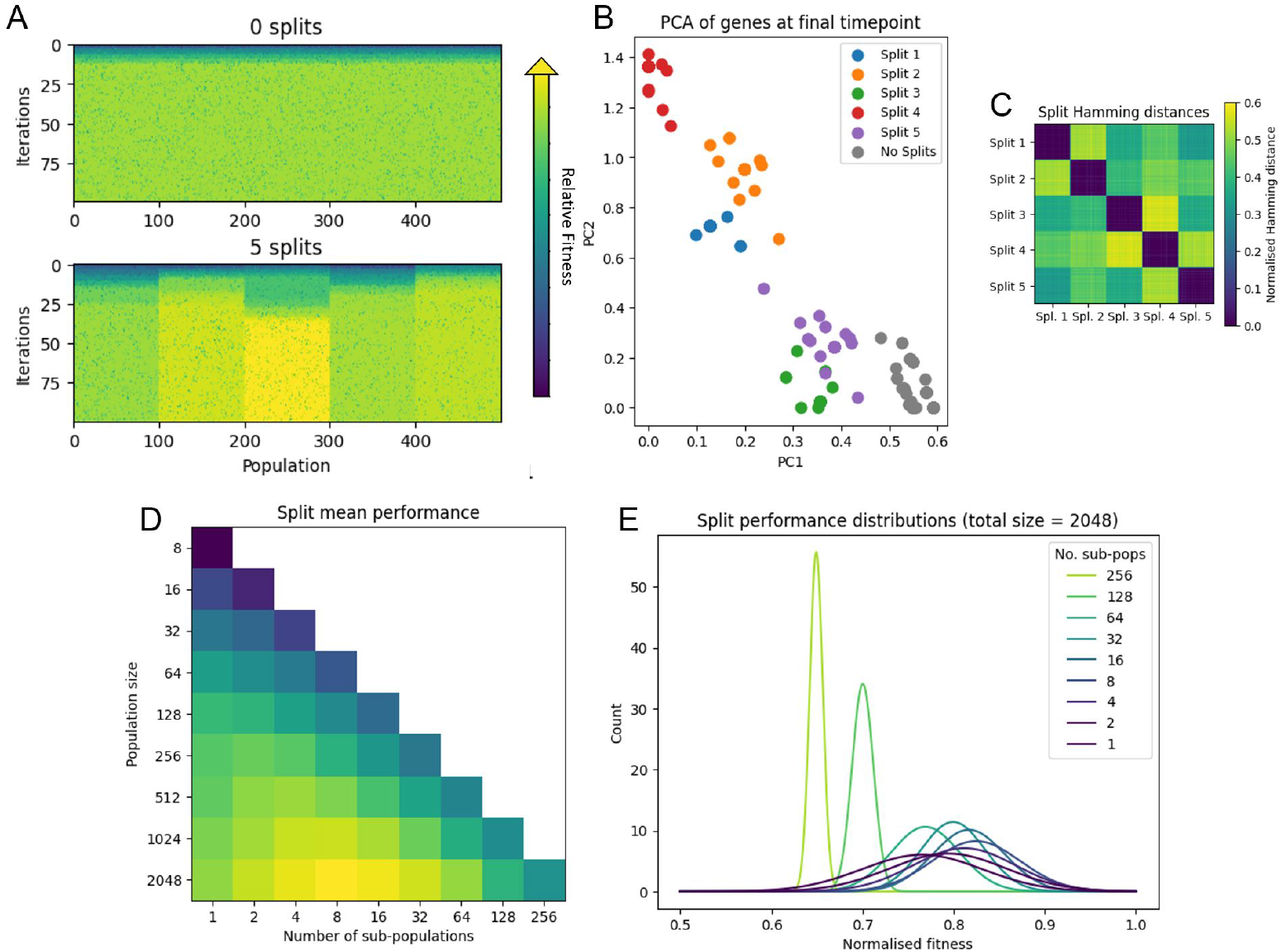
Investigating the effects of population splitting in directed evolution. A: Example of a directed evolution run split into five sub-populations vs a single large population. Colourbar indicates fitness value. B: Principal components analysis of the final time point sequences of a split vs non-split population. Hamming distances between the final timepoint sequences of C: a split population and D: a non-split population. E: Mean performance of directed evolution with varying total population and sub-population size. F: Distribution of performance with varying number of sub-populations (fit to a normal distribution). Experiments ran for 100 generations. N = 25, K = 5, mutations per cell = 0.1, selection threshold top 5%, as per [21].

Principal components analysis (PCA) was used to verify that sub-populations diverge on separate trajectories. The final timepoint gene sequences from the simulation in Fig.3A were collected. PCA was performed on the combined dataset to to reduce the 25-dimensional genetic space to just 2 dimensions for visualisation (Fig. 3B). The result shows that each sub-population forms a cluster, and the overall variation of the split population is significantly more than the non-split population. This is further verified in Fig. 3C, which displays that the average normalised Hamming distance between sub-populations (0.44) was far greater than that within sub-populations (0.011, similar to the average Hamming distance of 0.009 measured within the large single population).

Population splitting can clearly confer a benefit to performance, however there is a trade-off between splitting the population to maximise exploration, and keeping sub-populations large enough to effectively search around their local position on the landscape. Fig.3D summarises the performance of population splitting for different total population sizes, demonstrating that if the (total) population is too small, splitting is instead detrimental to performance. Fig.3E shows the distribution of performance outcomes, over 1000 runs, for splitting a large population into increasingly smaller populations. This demonstrates an additional advantage of population splitting, which is that the variance of the final outcome decreases as the number of sub-populations increases.

### Multi-dimensional selection with simulated novel selection methods

Previous sections have operated within the constraints of well-established directed evolution selection methods (e.g. FACS). Emerging methods for selection, however, may offer increased capabilities; notably using microfluidics to observe cells for long time periods prior to selection [16, 17]. Not only would long-term observation increase the reliability of readings and allow selection based on complex time-dependent traits, it would also allow for multiple properties (or responses to stimuli) to be measured in a single round of selection. Such multi-dimensional selection is highly applicable in the directed evolution of biosensors, in which one seeks to optimise both specificity and sensitivity [21].

The current standard approach to multi-dimensional selection (e.g. in FACS) is to perform sequential rounds of selection, one for each property [21]. This introduces a systematic error with respect to the selection objective, as shown in Fig. 4A. The blue dashed line divides the true best cells from the population (as could be achieved by single-round selection), whereas the orange dashed lines represent the cut-offs of a double-round selection setup. Cells that are poor in one property but excel in another are not selected by double-round selection. As a result, the overall performance of single-round selection is higher (Fig. 4B).

**Fig 4.**
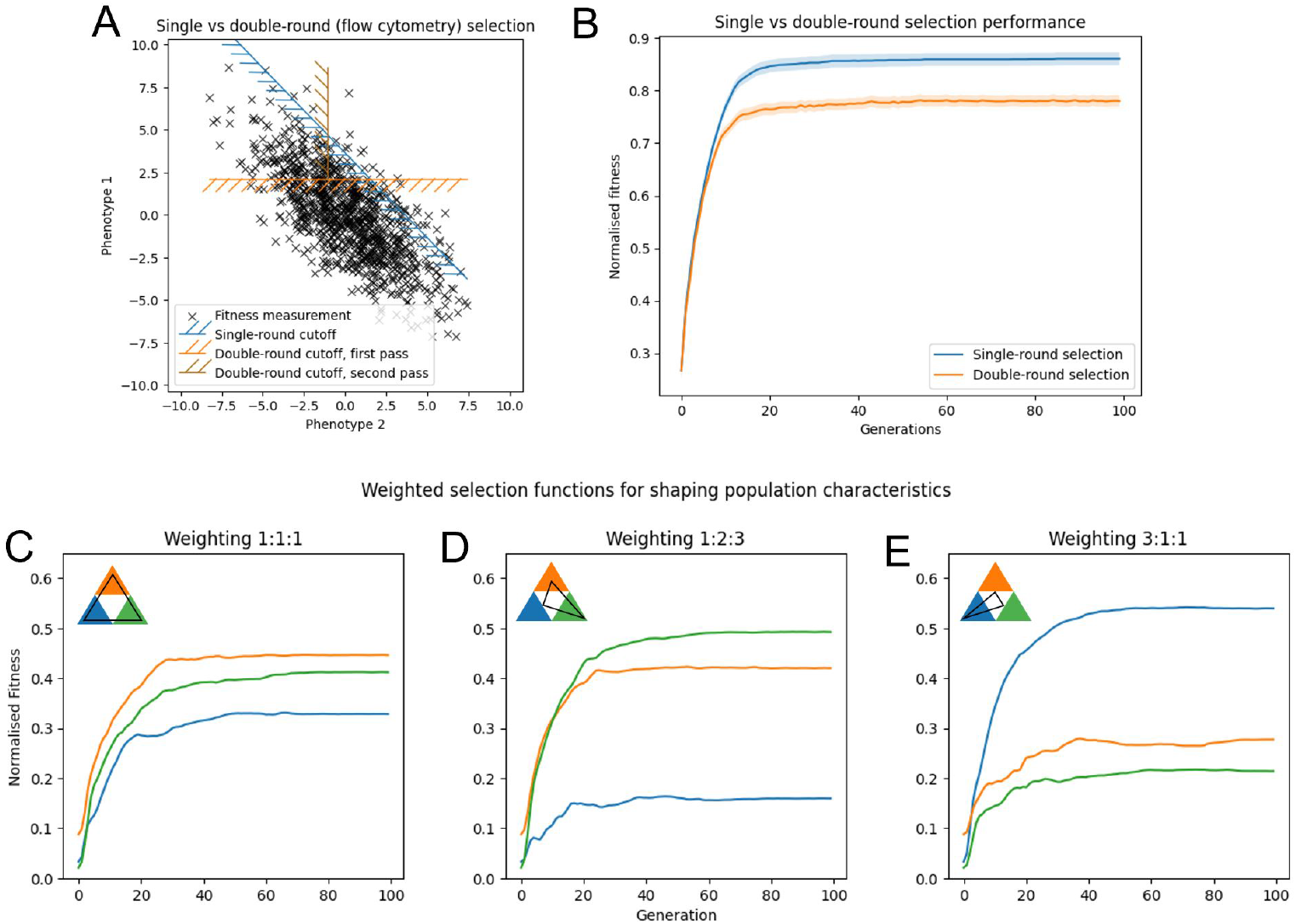
Optimisation of multi-dimensional selection. A: Demonstration of selection patterns with double-round (i.e. flow cytometry selection) and single-round selection. Randomly-generated fitness points normally-distributed around the mean 0 of phenotypes 1 and 2. B: Directed evolution performance of double-round vs single-round selection. Performance of weighted selection functions to perform directed evolution over three properties, with weightings of C: 1:1:1, D: 1:2:3 and E: 3:1:1 respectively. Experiments ran for 100 generations. N = 25, K = 5, mutations per cell = 0.1, population size = 1000, selection threshold top 5%, as per [21].

Not only would the described emerging methods improve upon FACS in the simplest case, they would also offer the additional ability to *tune* the prioritisation of different properties. When selecting on a single property, the fitness value (*F* ) used to determine selection is simply the value of that property. When selecting on multiple properties, however, the overall fitness value used to determine selection is some combination. Given that most directed evolution experiments have a limited amount of time and/or physical resources, it is crucial to consider how much one prioritises each property in this combination. This prioritisation can be implemented by applying a weight (*w*_*i*_) to the value of each property. So, *F* = *w*_1_*f*_1_ + *w*_2_*f*_2_ + *w*_3_*f*_3_, where *f*_*i*_ is the value of property *i*. In the simplest case, we allow all weightings to be equal (Fig. 4C). By changing the weightings of the properties we observe proportional gains in the fitness of each property (Fig. 4D and E).

### Translation to empirical fitness landscapes

The preceding results are based on simulations of NK landscapes, which although widely used, may not capture all important properties of real fitness landscapes. In order to test our strategies further, we therefore applied them to the empirical GB1 fitness landscape [19], which exhaustively measures the binding strength of 160,000 variants of the GB1 immunoglobulin protein to IgG-Fc.

We used this landscape to assess the performance of strategies that employ base chance and/or population splitting. Fig. 5A displays the performance of varying combinations of base chance and splitting. We observed a clear optimum in population splitting in the range of 20 sub-populations, which remains the optimum regardless of the base chance chosen. The trends with respect to base chance are much weaker, and also dependent on the number of sub-populations. In particular the optimal base chance decreases slightly from 0.2 to 0.1 as the number of sub-populations increases.

**Fig 5.**
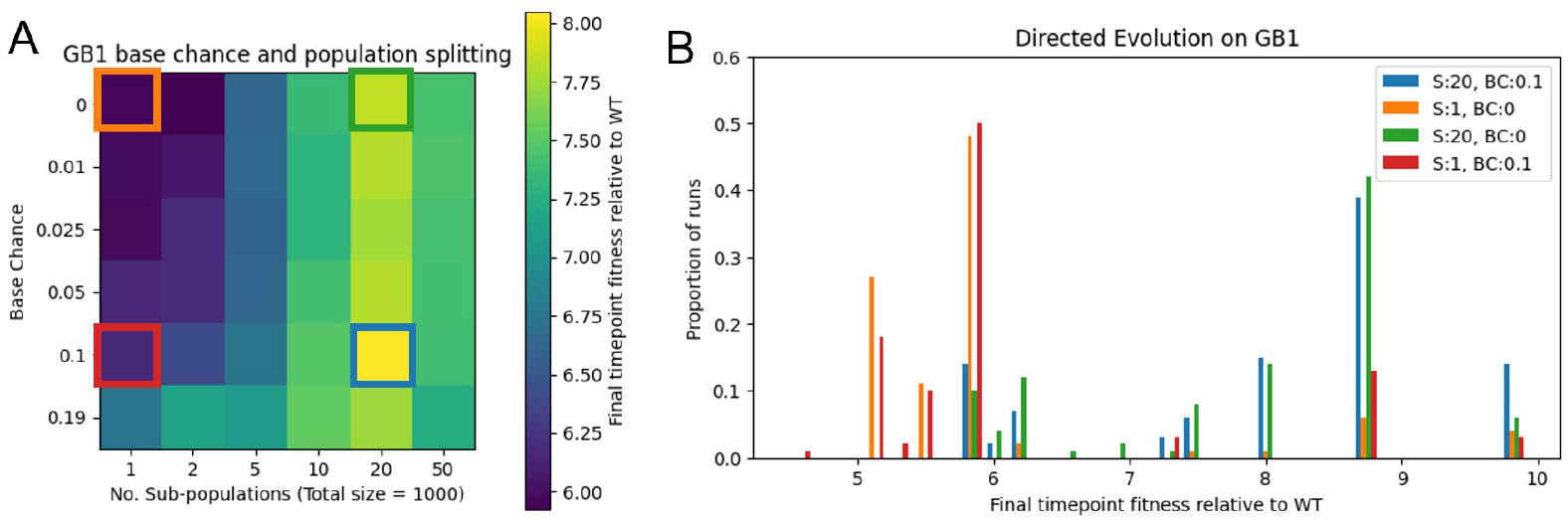
Application of selection function and population splitting to the empirical GB1 landscape. A: Performance of an array of base chance values and sub-population numbers in GB1 directed evolution. B: Distribution of outcomes on GB1 with varying values of base chance (BC) and sub-population numbers (S). Outcomes from 1000 simulations. GB1 max = 9.91, min = 0, wildtype simulation start = 1 (VDGV), mutation rate = 0.01, population size = 500.

Fig.5B displays the distribution of outcomes from GB1 directed evolution using four different strategies highlighted in Fig.5A (neither splitting nor base chance, each strategy in isolation, and both strategies combined). Of these four methods, the standard approach to directed evolution (employing neither splitting nor base chance) performs the worst, with over 50% of directed evolution runs on GB1 getting trapped at a local optimum (5.8x wildtype fitness). By introducing splitting, that frequency is reduced to less than 15%, and the mode outcome is 8.8x wildtype fitness. As for the fraction of runs which reach the *global optimum*, it is 10% with splitting, compared to less than 2% using the standard approach. Put another way, the best possible outcome of the experiment becomes *5 times more likely* with population splitting. By using both population splitting and base chance, this further increases to 15%, or 7.5 times more likely than the no splitting/base-chance case.

When deploying single-cell selection techniques in practice, it is not possible to identify the optimal parameters *before* beginning. However, considering Fig.5A and previous NK landscape results, we hypothesise that the standard approach of no splitting and no base chance may *in general* significantly under-perform in real-world experiments. Deviating from the standard approach to directed evolution, in particular by population splitting and/or adding a non-zero base chance, may therefore offer benefits even if it is not the absolute optimal strategy.

## Discussion

This study demonstrates that even in the absence of sequencing information, there are approaches that can be used to improve directed evolution outcomes. Such approaches were demonstrated both on simulated NK landscapes, and on the GB1 immunoglobulin protein empirical fitness landscape [19]. On the GB1 landscape, we showed that a population splitting strategy can lead to a five-fold increase in the probability of reaching the global optimum. This probability was shown to further increase by implementation of a “base chance” of selection, which aids in escaping local optima.

Given that empirical fitness landscapes, such as that of GB1, are scarce, the majority of this study relied on theoretical fitness models. These do not perfectly represent the statistical properties of natural fitness landscapes. In order to improve the reliability of simulations such as these, future work could tailor the NK model to the specific application of protein fitness landscapes. For instance, by allowing K to be drawn from a distribution, as opposed to a constant value for each amino acid position, or by integrating the information offered by PAM substitution matrices into fitness estimations. NK model variants also exist that emphasise the role of neutral drift, another factor that could be integrated into an alternative NK model [40, 41].

It was observed in this work that landscape structure (e.g. N and K values in the NK model) impacts the performance of directed evolution strategies. It would also be beneficial, therefore, to be able to infer landscape properties prior to performing an experiment. Our previous work demonstrated a method for inferring ruggedness using an FCN model trained to take in mutational spread data from a single starting location [42]. Similar landscape inference techniques have been performed using sequence alignment data, which assuming sufficient data exists online, also do not require sequencing during a directed evolution experiment [43].

With *de novo* protein design approaches still in their infancy, directed evolution remains an important component of the protein engineering toolbox. Here we described several approaches for directed evolution optimisation that do not require iterative sequencing or synthesis. Many of these approaches can be used with existing experimental techniques, and others are designed to support the advancements in single-cell selection technologies that we hope will improve the quality of directed evolution in future.

## Methods

### Model

To test our selection algorithms we implemented *in silico* simulations of directed evolution, which can be applied to either synthetic or empirical landscapes. We model each genetic variant within a population using a one-dimensional array of *N* integer values (Fig. 1), which is initialised at a random starting location (NK) or wildtype sequence (GB1), creating a population of size *P* . To calculate the corresponding fitness value of a gene, this array is either used as input to the NK model (Methods: NK Landscapes), or used as coordinates to look up fitness in an empirical data set (Methods: Empirical Landscapes). Our simulation algorithm proceeds with three cyclic steps; selection, proliferation, and mutation.

Individual cell fitness values are input to a selection function (Fig. 1), which translates relative fitness into probability of selection. Then, each cell is selected (or not) based only on its respective probability of selection. Cells that are selected are proliferated to bring the population back up to its original size. To perform proliferation, a *new* population of size *P* is created by randomly sampling (with replacement) from the previously selected cells. This introduces a degree of stochasticity mirroring experimental error and biological variation, as described in other models of evolutionary processes [44].

Once a population of size *P* has been selected and proliferated, mutations are introduced by making random changes to genes in the population. For every cell’s genetic code (an array), each residue (i.e. nucleotide or amino acid) has a random chance, *p*_*I*_, of changing into another random *different* value (i.e., each other possibility is equally likely). Given the gene length *N*, the expected number of changes across the entire gene (the “mutations per cell”) is given by *µ* = *Np*_*I*_ . This quantity is useful to work with, as a fixed *µ* will give comparable results as *N* varies.

The final result is a new population of size *P*, and the process of selection, proliferation, and mutation may be repeated again. In this work, it is assumed that this runs for a fixed number of iterations, and the final result (by which we compare methods) is the maximum fitness across the final population. ‘Normalised fitness’ divides this measure by the global maximum of the landscape.

### Selection Functions

In this work, we introduce the concept of a “selection fuction”. This function takes a cell’s fitness percentile, defined as the percentage of other cells in the population it is fitter than, and outputs its chance of being selected to be in the next population. The selection function used in this work is defined by two parameters: a “threshold” and a “base chance”(Fig. 2A). Above the “threshold” the selection function outputs 1, otherwise it outputs the “base chance” (Equation 1). In this work, selection functions are normalised to select a constant fraction of cells (20% throughout this paper). This is to ensure that in a real continuous directed evolution experiment, the time required for proliferating cells between iterations remains effectively constant such that a “fair” comparison is made between strategies in terms of performance per-unit-time. By fixing the selected proportion (given by the integral of the selection function), our parameter search space is reduced to one dimension, as every base chance has only one possible corresponding threshold value (Equation 2).

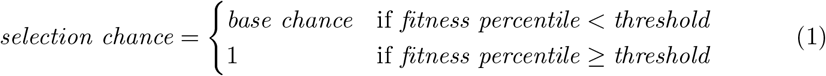

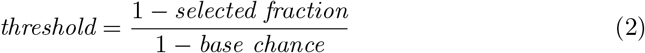

### NK Landscapes

The NK model is a widely used approach for generating synthetic fitness landscapes with tuneable ruggedness [33, 34]. In the NK model, a gene is represented by an array of *N* sites, each of which has *A* possible values. In this work we will generally set *A* = 2 such that each entry has two possibilities (i.e. binary 1 or 0). Every entry corresponds to a “locus”, which interacts with *K* other loci in the gene. The fitness contribution of each locus is dependent on the state of that locus and the state of the *K* other loci it interacts with . The fitness *F* of a gene *G* is the sum of the fitnesses of each locus. When *K* = 0, all loci are independent and the model is linear (and hence has a single peak). The other extreme, *K* = *N* − 1, in which every locus interacts with every other locus, is maximally unstructured; the fitness landscape consisting of only random noise.

The NK model defines a parameterised *distribution* over functions *F* : *A*^*N*^ *→* ℝ. To define how to generate samples from this distribution, first let *L* : *N × K → N* be a “locus function”, where *L*(*a, i*) is the *i*th site that locus *a* interacts with. We require each locus to interact with precisely *K* other, uniformly random, sites, independent of all other loci. We then also have *N · A*^*K*+1^ independent, identically distributed standard normal (𝒩 (0, 1)) random variables, which we denote as 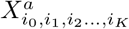, where *a ∈* {1…*N* } and *i*_0,1,…,*K*_ *∈* {1…*A*}. These represent the possible fitness contributions of each loci, with *a* being the index of the loci in question, and *i*_0,1,…,*K*_ being the values of that loci and the *K* other loci it interacts with, plus itself (*i*_0_).

Then, *F* is given as the following, where *g ∈ A*^*N*^, and *g*[*i*] denotes the *i*th site in *g*:

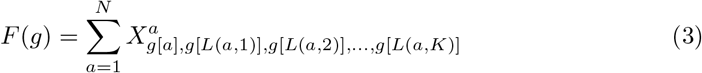

Computationally, the algorithm used explicitly generates and stores *L*; in particular as a *N × N* binary matrix. However, it does not store the *N · A*^*K*+1^ random variables explicitly as doing so would require far too much memory. Instead, every time the value of 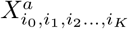 is required, *a, i*_0_, *i*_1_, …, *i*_*K*_ are used as inputs to a pre-defined deterministic pseudorandom generation algorithm. As a result, to “look up” the value of 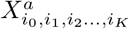 the same computation process is repeated each time.

### Empirical Landscapes

Empirical fitness landscapes are real data sets produced by measuring the fitness of all possible sequential variants of a protein (or region of a protein). The empirical landscape used in this paper is that of GB1, an immunoglobulin-binding protein found in Streptococcal bacteria. The fitness of each GB1 variant was determined by stability and binding to IgG-Fc [19]. For our algorithm, the landscape is stored as an four-dimensional array, where each dimension corresponds to a variable residue. By assigning every amino acid a number from 1 to 20, each sequence of amino acids can therefore be mapped to a set of coordinates that points to the corresponding fitness value in the array. The landscape is hence equivalent to *N* = 4 and *A* = 20 with the parameters defined above in the NK Landscapes section. All directed evolution simulations were started from the wild-type sequence “VDGV”.

## Acknowledgments

H.S. is supported in part by the Engineering and Physical Sciences Research Council (EPSRC) projects EP/W000326/1 and EP/X017982/1.

## Data Availability

All data and code used for running experiments and plotting is available on a GitHub repository at https://github.com/nesou2/direvo_sim.git. Code is also available on the Software Heritage Archive: swh:1:dir:e524b6ffc471276a6b0cf001604655d9d5fbc0f0.

## References

1. Yan X, Liu X, Zhao C, Chen G. Applications of synthetic biology in medical and pharmaceutical fields. Signal Transduct Target Ther. 2023;8:199. doi:10.1038/s41392-023-01440-5.

2. Scown CD, Keasling JD. Sustainable manufacturing with synthetic biology. Nature Biotechnology. 2022;40(3):304–307. doi:10.1038/s41587-022-01248-8.

3. Sargent D, Conaty WC, Tissue DT, Sharwood RE. Synthetic biology and opportunities within agricultural crops. Journal of Sustainable Agriculture and Environment. 2022;1(2):89–107. doi:10.1002/sae2.12014.

4. Dauparas J, et al. Robust deep learning–based protein sequence design using ProteinMPNN. Science. 2022;378(6615):49–56. doi:10.1126/science.add2187.

5. Ferruz N, Schmidt S, Höcker B. ProtGPT2 is a deep unsupervised language model for protein design. Nature Communications. 2022;13(1):4348. doi:10.1038/s41467-022-32007-7.

6. Singer JM, et al. Large-scale design and refinement of stable proteins using sequence-only models. PLOS ONE. 2022;17(3):e0265020. doi:10.1371/journal.pone.0265020.

7. Anishchenko I, et al. De novo protein design by deep network hallucination. Nature. 2021;600(7889):547–552. doi:10.1038/s41586-021-04184-w.

8. Wicky BIM, et al. Hallucinating symmetric protein assemblies. Science. 2022;378(6615):56–61. doi:10.1126/science.add1964.

9. Arnold FH. Design by Directed Evolution. Accounts of Chemical Research. 1998;31(3):125–131. doi:10.1021/ar960017f.

10. Nixon AE, Sexton DJ, Ladner RC. Drugs derived from phage display. mAbs. 2014;6(1):73–85. doi:10.4161/mabs.27240.

11. Heater BS, et al. Directed evolution of a genetically encoded immobilized lipase for the efficient production of biodiesel from waste cooking oil. Biotechnology for Biofuels. 2019;12(1):165. doi:10.1186/s13068-019-1509-5.

12. Neuenschwander M, Butz M, Heintz C, Kast P, Hilvert D. A simple selection strategy for evolving highly efficient enzymes Nat Biotechnol. 2007;25(10):1145–1147. doi:10.1038/nbt1341.

13. Wu Y, Jameel A, Xing X, Zhang C. Advanced strategies and tools to facilitate and streamline microbial adaptive laboratory evolution Trends in Biotechnology. 2022;41(1):38–59. doi:10.1016/j.tibtech.2021.04.002.

14. Yang G, Withers SG. Ultrahigh-throughput FACS-based screening for directed enzyme evolution Chembiochem. 2009;10(17):2704–2715. doi:10.1002/cbic.200900384.

15. Trivedi VD, Mohan K, Chappel TC, Mays ZJS, Nair NU. Cheating the cheater: Suppressing false positive enrichment during biosensor-guided biocatalyst engineering ACS Synth Biol. 2022;11(1):420–429. doi:10.1021/acssynbio.1c00506.

16. Luro S, Potvin-Trottier L, Okumus B, Paulsson J. Isolating live cells after high-throughput, long-term, time-lapse microscopy Nat Methods. 2020;17(1). doi:10.1038/s41592-019-0620-7.

17. Sheets MB, Tague N, Dunlop MJ. An Optogenetic Toolkit for Light-Inducible Antibiotic Resistance Nat Commun. 2023;14(1). doi:10.1038/s41467-023-36670-2.

18. Wright S. The Roles of Mutation, Inbreeding, crossbreeding and Selection in Evolution. Proceedings of the XI International Congress of Genetics. 1932;8:209–222.

19. Wu NC, et al. Adaptation in protein fitness landscapes is facilitated by indirect paths. eLife. 2016;5:e16965. doi:10.7554/eLife.16965.

20. Papkou A, et al. A rugged yet easily navigable fitness landscape. Science. 2023;382(6673):eadh3860. doi:10.1126/science.adh3860.

21. Leopoldo FM, et al. Directed evolution of the PcaV allosteric transcription factor to generate a biosensor for aromatic aldehydes. Journal of Biological Engineering. 2019;13(1):91. doi:10.1186/s13036-019-0214-z.

22. LaCroix RA, Palsson BO, Feist AM. A Model for Designing Adaptive Laboratory Evolution Experiments. Applied and Environmental Microbiology. 2017;83(8):e03115–16. doi:10.1128/AEM.03115-16.

23. Romero PA, Arnold FH. Exploring protein fitness landscapes by directed evolution. Nature Reviews Molecular Cell Biology. 2009;10(12):866–876. doi:10.1038/nrm2805.

24. Fox R, et al. Optimizing the search algorithm for protein engineering by directed evolution. Protein Engineering, Design and Selection. 2003;16(8):589–597. doi:10.1093/protein/gzg077.

25. Wu Z, et al. Machine learning-assisted directed protein evolution with combinatorial libraries. Proceedings of the National Academy of Sciences. 2019;116(18):8852–8858. doi:10.1073/pnas.1901979116.

26. Wittmann BJ, Yue Y, Arnold FH. Informed training set design enables efficient machine learning-assisted directed protein evolution. Cell Systems. 2021;12(11):1026–1045.e7. doi:10.1016/j.cels.2021.07.008.

27. Yang KK, Wu Z, Arnold FH. Machine-learning-guided directed evolution for protein engineering. Nature Methods. 2019;16(8):687–694. doi:10.1038/s41592-019-0496-6.

28. Frisby TS, Langmead CJ. Bayesian optimization with evolutionary and structure-based regularization for directed protein evolution. Algorithms for Molecular Biology. 2021;16(1):13. doi:10.1186/s13015-021-00195-4.

29. Hu R, et al. Protein engineering via Bayesian optimization-guided evolutionary algorithm and robotic experiments. Briefings in Bioinformatics. 2023;24(1). doi:10.1093/bib/bbac570.

30. D’Costa S, et al. Inferring protein fitness landscapes from laboratory evolution experiments. PLOS Computational Biology. 2023;19(3):e1010956. doi:10.1371/journal.pcbi.1010956.

31. Molina RS, et al. In vivo hypermutation and continuous evolution. Nature Reviews Methods Primers. 2022;2(1):1–22. doi:10.1038/s43586-022-00119-5.

32. Carpenter AC, et al. Have you tried turning it off and on again? Oscillating selection to enhance fitness-landscape traversal in adaptive laboratory evolution experiments. Metabolic Engineering Communications. 2023;17:e00227. doi:10.1016/j.mec.2023.e00227

33. Kauffman S, Levin S. Towards a general theory of adaptive walks on rugged landscapes Journal of Theoretical Biology. 1987;128(1):11–45

34. Kauffman S, Weinberger ED. The NK model of rugged fitness landscapes and its application to maturation of the immune response Journal of Theoretical Biology. 1989;141(2):211–245

35. Iwasa Y, Franziska M, Nowak MA. Stochastic tunnels in evolutionary dynamics. Genetics. 2004;166(3):1571–1579 doi:10.1534/genetics.166.3.1571

36. Ochs IE, Desai MM. The competition between simple and complex evolutionary trajectories in asexual populations BMC Evolutionary Biology. 2015;15(1) doi:10.1186/s12862-015-0334-0

37. Vedal LS, Isalan M, Heap JT, Ledesma-Amaro R. A primer to directed evolution: current methodologies and future directions Royal Society of Chemistry. 2023. doi: DOI10.1039/D2CB00231K

38. Gavrilets, S. Perspective: Models of Speciation: What Have We Learned in 40 Years? Society for the Study of Evolution, Wiley. 2003;57(10):2197–2215. 10.1111/j.0014-3820.2003.tb00233.x

39. Chen Z, et al. Multi-Population Evolutionary Algorithm for Solving Constrained Optimization Problems Artificial Intelligence Applications and Innovations (IFIP Conference). 2005.

40. Barnett L. Ruggedness and neutrality—the NKp family of fitness landscapes ALIFE: Proceedings of the sixth international conference on Artificial life. 1998. pp18–27.

41. Newman MEJ, Engelhardt R. Effects of neutral selection on the evolution of molecular species 1997.

42. Towers S, James J, Steel H, Kempf I. Learning-Based Estimation of Fitness Landscape Ruggedness for Directed Evolution BioRxiv. 2024 doi:10.1101/2024.02.28.582468.

43. Thomas N, Agarwala A, Belanger D, Song YS, Colwell LJ. Tuned Fitness Landscapes for Benchmarking Model-Guided Protein Design BioRxiv. 2022. 10.1101/2022.10.28.514293

44. Desai MM, Fisher DS. Beneficial Mutation-Selection Balance and the Effect of Linkage on Positive Selection Genetics. 2007;176(1):1759–1798. 10.1534/genetics.106.067678

